# Gaining insight into the allometric scaling of trees by utilizing 3d reconstructed tree models - a SimpleForest study

**DOI:** 10.1101/2022.05.05.490069

**Authors:** Jan Hackenberg, Mathias Disney, Jean-Daniel Bontemps

## Abstract

Forestry utilizes volume predictor functions utilizing as input the diameter at breast height. Some of those functions take the power form *Y* = *a* ∗ *X*^*b*^. In fact this function is fundamental for the biology field of allometric scaling theories founded round about a century ago. The theory describes the relationships between organs/body parts and the complete body of organisms.

With digital methods we can generate 3d forest point clouds non destructively in short time frames. SimpleForest is one free available tool which generates fully automated ground and tree models from high resoluted forest plots. Generated topological ordered cylinder models are called commonly QSMs.

We use SimpleForest QSMs an build a function which estimates the total supported wood volume at any given point of the tree. As input we use the supported soft wood volume for those query points. Instead of measuring directly the soft wood volume we use as a proxy the number of supported twigs. We argue with the pipe model theory for the correctness of the proxy.

We can use the named relationship to also filter our QSMs made of an open data set of tree clouds. The filter corrects overestimated radii. And we compare the corrected QSM volume against the harvested reference data for 66 felled trees. We also found QSM data of TreeQSM, a competitive and broadly accepted QSM modeling tool. Our RMSE was less than 40% of the tree QSM RMSE. And for other error measures, the r^2^_adj_. and the CCC, the relative improvement looked even better with 27% and 21% respectively.

We consider this manuscript as highly impactful because of the magnitude of quality improvement we do. The relation between soft volume and total volume distributions seems to be really strong and tree data can easily also be used as example data for the generic field of allometric scaling.

## 1 INTRODUCTION

### 1.1 Allometric Theory

Many ways to capture the bio volume of trees with mathematical concepts have been performed over the last centuries. Already 500 years ago Leonardo da Vinci noticed that the sum of cross sectional areas at any height equals the cross sectional area of the trunc (Richter et al., 1970). Additionally in the named work the comparison of a tree’s branching system with a river network was performed.

Pressler’s law (Pressler and Pressler, 1865) refined the theory for deciduous temperate species. As new leaves have to be fed with nutrients via new pipes the summed up cross sectional areas of pipes formed in the latest growing season correlates to the annual diameter increment of the trunc.

The latter theory was then even further generalized by accounting to inactive pipes in the so called Pipe Model Theory (PMT) (Shinozaki et al., 1964). Due to thinning and self thinning pipes become inactive and build up the heartwood. Those inactive pipes are located in the center of the stem or branching units. The active pipes which feed the leaves or set of leaves with nutrient build up the outer perimeter of the wooden structure. We can call this outer part sapwood. We can see the whole theory abstracted in figure 1.

**FIGURE 1.**
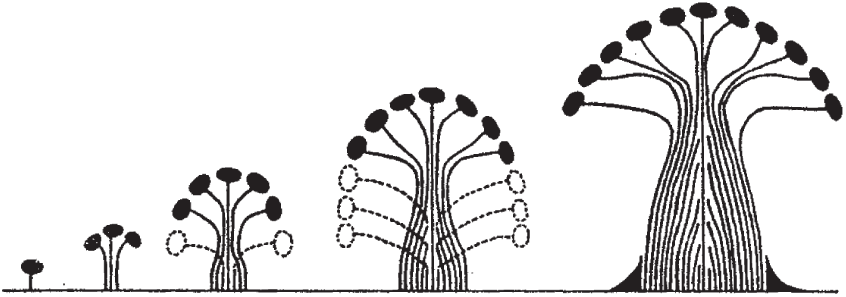
The original figure used to publish the PMT (Shinozaki et al., 1964). The leaves depicted in black dots are fed by active pipes in the outer sapwood. The inner pipes are inactive as the leaves are pruned with their pipe endings. Those inactive pipes build the inner heartwood.

Recently a well written summary about the PMT and related tree describing theories was published (Lehnebach et al., 2018). We get insight how various theories explain tree properties, see figure 2 for an abstract view which refers also to the previous mentioned Pressler’s law and the Da Vinci’s rule.

**FIGURE 2.**
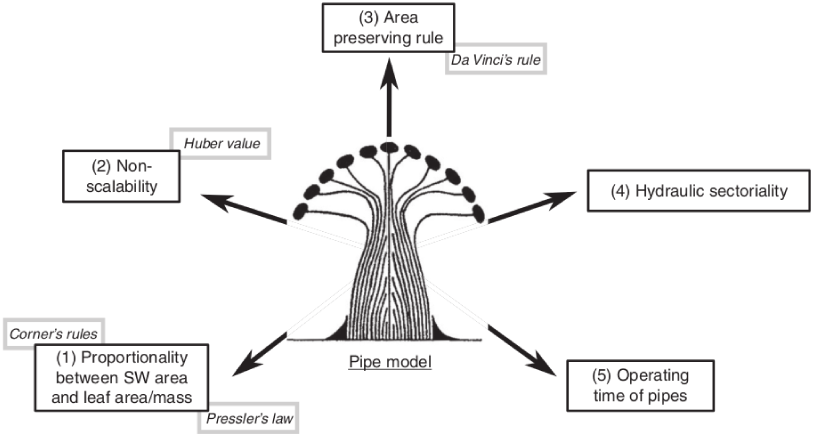
The original figure used to publish in a review paper about the PMT (Lehnebach et al., 2018). We see five different tree properties linked to explanatory theories.

In fact the PMT can be seen as a specialized case of the allometric scaling theory. Allometric scaling is the biology field where mathematical relationships between organs/body parts and the complete body of organisms are build. A century ago the scaling of shore-crab bodies was described to follow a pattern which can be described with the two parameter power function seen in equation 1 (Huxley, 1924).

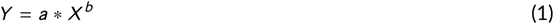

Up to day allometric scaling theory is a relevant field and has been refined throughout the decades (Packard, 2020). It was found meaningful also in the development of the West, Brown and Enquist (WBE) model in forest ecology (West et al., 2009; Enquist et al., 2009). Astonishingly, the publication of the WBE model relies on more than fifty year old data which was presented with the publishing of the PMT (Shinozaki et al., 1964) in the sixties of last century. Both works show a significant impact and yet rely on few data.

### 1.2 Tree data availability for allometric scaling research

In forest ecology as well as in forestry when traditional measurement techniques are applied researchers often rely on only a few single tree parameter measures. The Diameter at Breast Height (`DBH`), the tree height (`H`) and the species category are recorded within national forest inventories (NFIs). The accuracy of those measurements is effected by a human bias. From the named parameters which are easy to obtain the volume of above ground biomass (Volume_AGB_) can then be estimated via allometric functions (Picard et al., 2015). An exemplary exponential model is show in equation 2 with *a* and *b* being the regression coefficients.

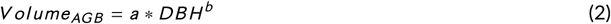

This model is of the from we already saw in equation 1. Yet such functions are costly to generate with traditional methods as here the felling and weighing of the tree is mandatory.

Equation 3 (van Laar and Akça, 2007) is another way to compute the Volume with the tree species specific from factor *f* describing the shape and taper of the trunc.

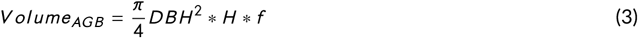

We can find those and other similar tree species specific equations in online data bases (Henry et al., 2013; Zianis et al., 2005). Nevertheless, utilizing those equations comes with a large uncertainty about the prediction accuracy, especially if regional growth pattern are accounted to. And as with traditional methods the Volume_AGB_ could only be destructively collected with high costs such functions and tables were not often published.

By increasing the field costs additionally more diameters at defined heights can be collected to build taper functions from. And with destructive techniques even the diameter of branches of various branch orders can be collected. But this is rarely performed due to time constraints in addition to other costs and available inventory data is limited.

Within the field of computational forestry the 3d reconstruction of high resoluted tree point clouds recorded by Light Detection and Ranging (LiDaR) sensors to serve as an alternative data collection tool became more and more attractive. More measurements without human bias are possible to perform in a much shorter time frame. The tree surface can easily be covered with multiple millions of euclidean points when high resoluted terrestrial laser scans (TLS) are recorded.

Since around a decade various software tools to produce so called Quantitative Structure Models (QSMs) have been published. QSMs consist of thousands and sometimes even ten or hundred thousands geometrical building bricks. Commonly the cylinder is utilized as the building brick (Markku et al., 2015). With the complete branching structure being geometrically captured the Volume_AGB_ can be measured directly and the human bias during data collection is minimized. Data collection time is roughly in the time frame of few hours or days when collecting TLS data which covers up to multiple hectares of forest. The mentioned data bases (Henry et al., 2013; Zianis et al., 2005) can therefore theoretically be filled with a magnitude of new functions even accounting to growth patterns, species provenance and site variability (Hackenberg et al., 2014). Next to volume `DBH` and `H` can be directly extracted from QSMs in their table formatted text output forms.

In addition to the geometrical description also topology information is stored inside a QSM. Figure 3 shows a real QSM of a complex tree’s point cloud with more than 35 thousand cylinders. And in figure 4 we draw an idealized and simplified mathematical QSM. We discuss the topological order represented by the informatics tree structure later to create new relevant tree parameters.

**FIGURE 3.**
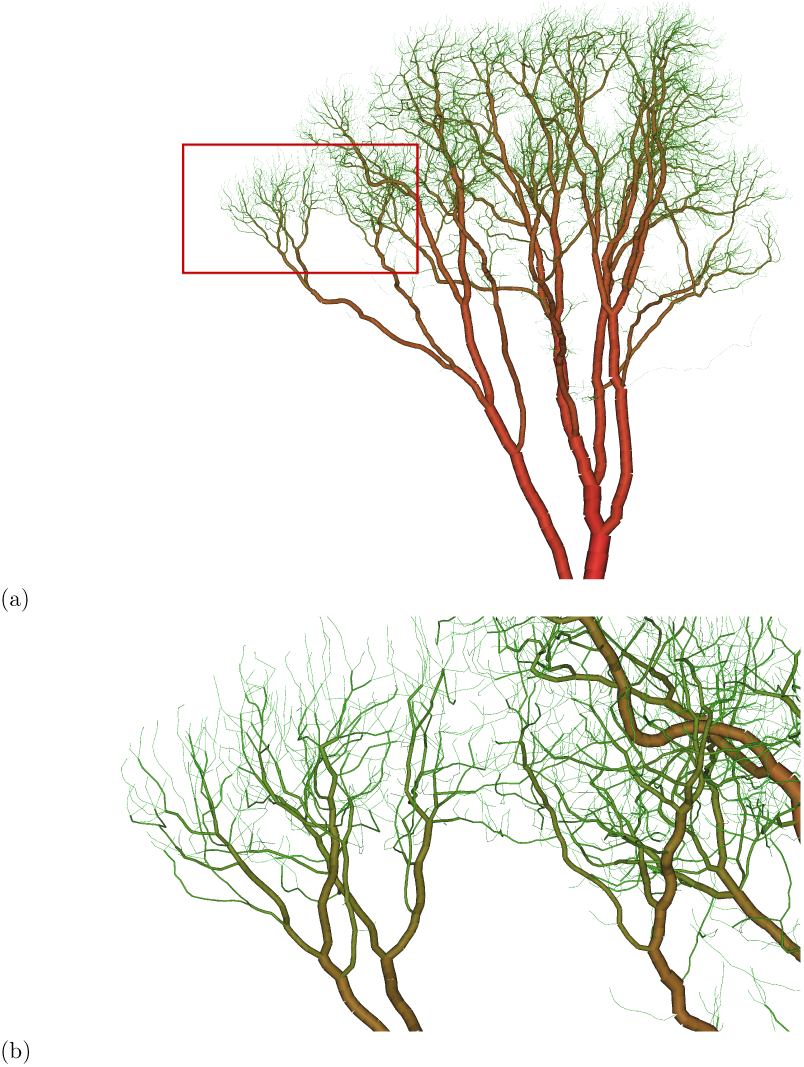
(a) A QSM of an *Acer pseudoplatanus* (Disney et al., 2018; Calders et al., 2018) with a DBH of ∼ 1.4 *m* containing approximately 25 *m*^3^ of woody biomass. The tree is located in Wytham Woods, UK. The QSM consists of more than 35 000 cylinders. (b) An enlarged screenshot of the area highlighted by the red bounding box in (a).

**FIGURE 4.**
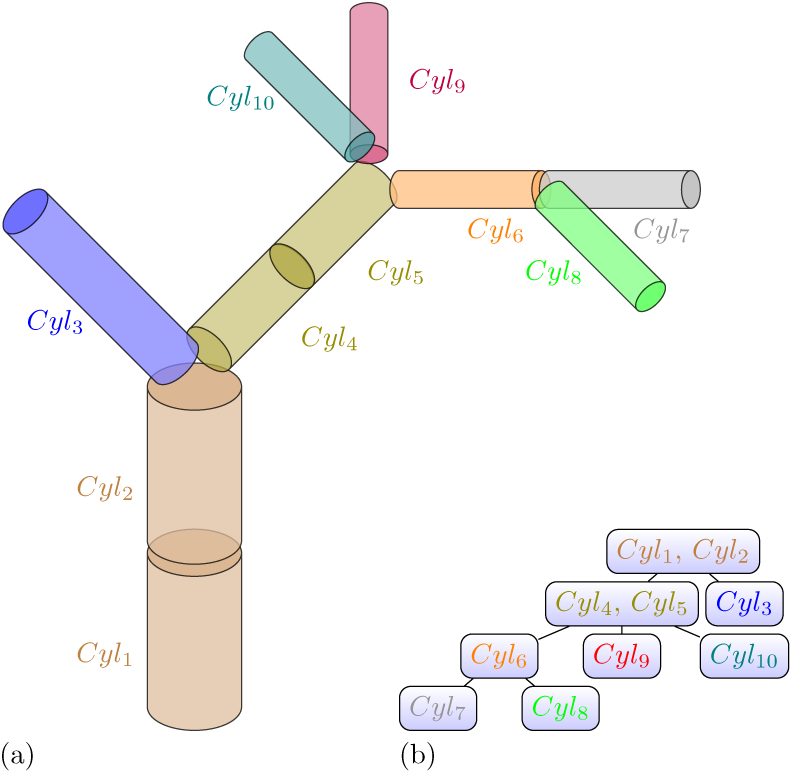
(a) An artificial QSM consisting of ten cylinders Cyl_n_. Each cylinder consists of two 3d points `start` and `end` as well as the `radius`. The cylinders between two neighboring branching junctions are combined to a list structure called segment. Each segment is colored differently. The cylinder list is always ordered from closest to the root to farthest away from the root. (b) We can see the topological order information from the QSM in (c). The segments are stored in the nodes. The root node is the segment which contains the root cylinder. The child nodes are always ordered by the size of the subtree.

Licensed under copy left SimpleTree (Hackenberg et al., 2014, 2015a), TreeQSM (Raumonen et al., 2013, 2015) or 3dForest (Trochta et al., 2017) are QSM producing tools which can be downloaded with their source code being free.

The free tools have been tested with by comparing QSM estimated Volume_AGB_ derived from TLS data versus destructive collected Volume_AGB_ field measurements (Hackenberg et al., 2021). But there has also been a lot of progress in utilizing other LiDar technologies such as airborne laser scanning (Kankare et al., 2013). We will use QSMs created with SimpleForest in the following study.

Additionally, we need to be aware that low cost photogrammetric sensors can serve as data collection tools when utilizing convolutional neural networks (Russell and Norvig, 2009) to compute depth maps from images. Those can be transformed to 3d point clouds which then can be utilized as input data for SimpleForest, TreeQSM, 3dForest or other tools. Constant break throughs (Chang and Chen, 2018; Tankovich et al., 2021; Lipson et al., 2021; Li et al., 2022) regarding accuracy of the derived photogrammetric clouds have been made by utilizing stereo vision cameras, which cost only about one tenth of the investment needed to purchase a TLS device.

As nowadays researchers without access to a smart phone camera are rare, the fact that even with every day photogrammetric sensors we can retrieve accurate point cloud representations is of huge impact. The clouds can be calculated from multiple un-referenced images of various point of views inside a scenery (Kuhn et al., 2020; Zhang et al., 2020). Even monocular methods for producing highly accurate depth maps from single photographies exist(Yuan et al., 2022; Alhashim and Wonka, 2018).

Additionally it is worth mentioning that future superior algorithms will produce more accurate data compared to nowadays algorithms on the same input data. We live in a time where we can create data bases with millions or even billions of tree measurements with every day technology. QSMs will be produced with free available tools on photogrammetric clouds and here we will look onto a few capabilities of QSMs to make progress towards finding answer for questions in the field of allometric scaling theory.

## 2 METHODS

### 2.1 Branch Order

Traditionally, the `Branch Order` (`BO`) is counted from the stem up to the tips. In branch order one, not only the largest branch segment is included, but also medium sized branches and even epicormic shoots. Therefore the correlation between this branch order and the diameter of segments of a certain branch order is rather low. To define a more meaningful order in the sense of diameter scaling we can read the inverse branch order from a tree.

### 2.2 Reverse Branch Order

#### Reverse Branch Order

The `Reverse Branch Order` (`RBO`) of a cylinder is the maximum depth of the subtree of the segment’s node. If we look at the real tree, the `RBO` denotes the maximal number of branching splits of the sub-branch growing out the segment.

We can see the inverse branch order depicted in the schematic QSM visible in figure 5.

**FIGURE 5.**
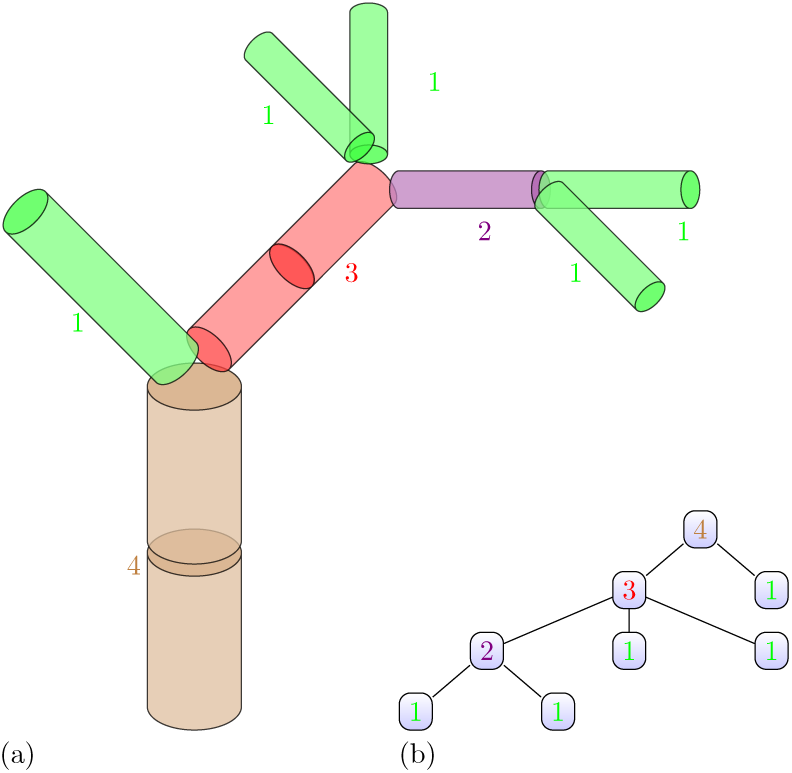
(a) The artificial QSM of figure 3 labeled with the `RBO`. (b) We can see the topological order information from the QSM in (a) and each node has the `RBO` displayed.

We can see a stronger correlation for the `RBO` compared to the `BO` if we look into the scatter plots of `BO` vs `radius` and `RBO` vs `radius`. Those plots are embedded in figure 6 (a) + (b).

**FIGURE 6.**
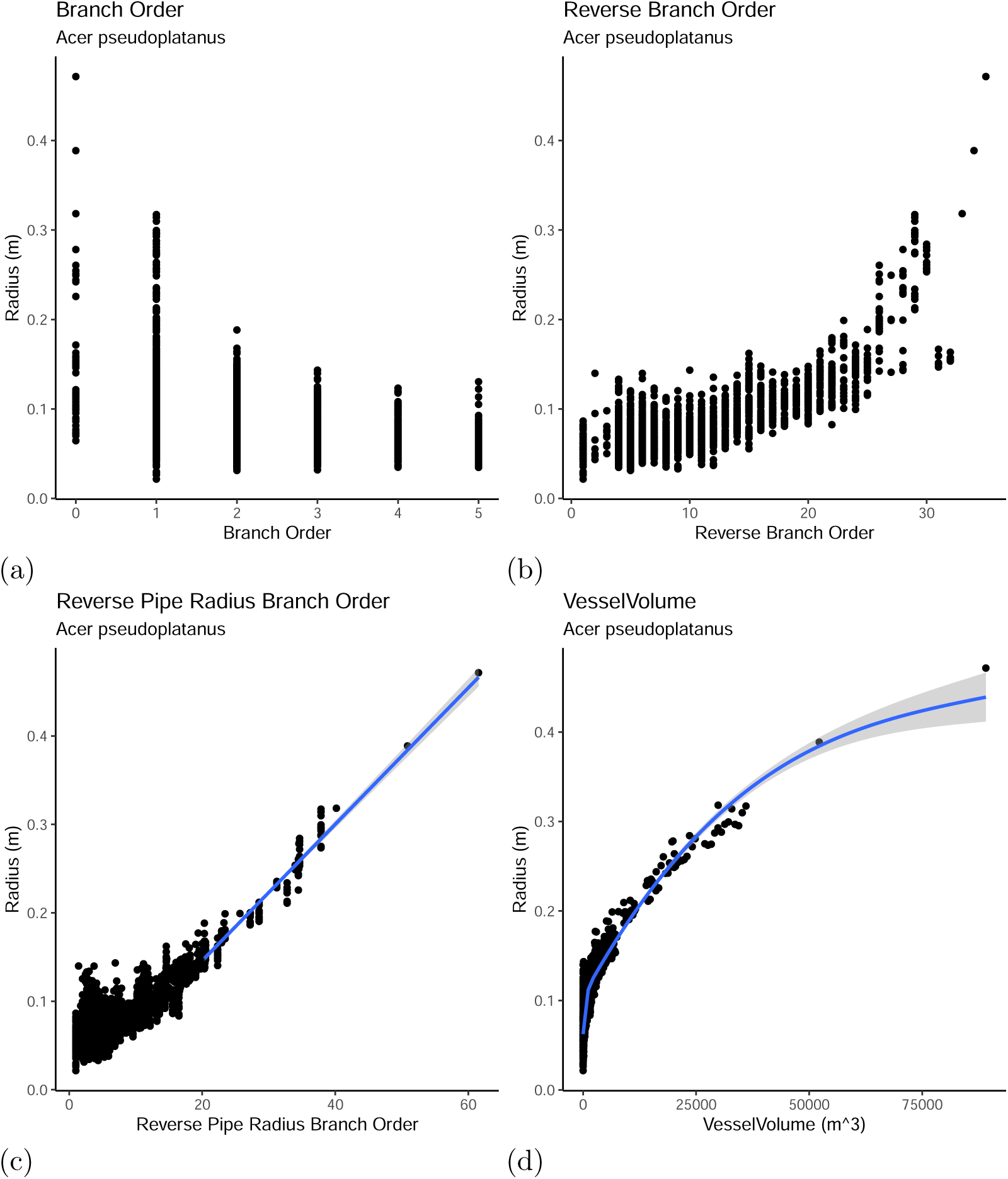
(a) We see the correlation between the traditional `BO` and the `radius` is not strong. Nevertheless the maximum value per branch order reveals a parabolic pattern. (b) We see the correlation between the `RBO` and the `radius` is stronger than the one depicted in (a). We see a noisy non linear pattern. (c) When looking at the scatter plot of the `RPRBO` vs the `radius` we can see a linear pattern, less noisy than the pattern in (b). (d) A power function form non linear pattern is visible when we plot the `VesselVolume` vs the `radius`. This pattern is the best in the sense that we barely see noise here.

#### Application for the Reverse Branch Order

We are about to define two useful predictors which can be read from a QSM to predict the radius. No matter if linear or non linear fitting techniques are utilized it is a good behavior to prepare the QSM data. As overfitting during the QSM generation is more likely to happen at the twig regions (Demol et al., 2022), we can simply create one filter to get rid of noisy cylinders by thresholding the `RBO`. For example Requesting the the `RBO` of valid cylinders has to be larger one removes all tip segments.

The remaining cylinders should then pass a second filtering routine. We recommend statistical outlier filtering utilizing the cloud to cylinder distance. Yet we don’t discuss details here, as this second filter is a point cloud based filter and does not fit the scope.

### 2.3 Reverse Pipe Area Branch Order

#### Reverse Pipe Area Branch Order

Firstly we define the Reverse Pipe Area Branchorder (`RPABO`): The WBE model considers that the branching architecture is self-similar (Brummer et al., 2017). In a self-similar branching architecture the `radius` as well as the cross sectional area of a twig is constant. By another definition *RPABO* is a tree specific area unit which equals one for a tip cylinder. We can always set it one as we argue this measure is of an unknown metrical unit. Knowledge of the real unit is unnecessary as the factor is crossed out during later calculations. Then by following the pipe model theory (Shinozaki et al., 1964) the *RPABO* of a cylinder at any location inside the QSM equals the number of supported tips. Supported tip cylinders always have a recursively defined parent relation to the query cylinder.

We can see the `RPABO` depicted in the schematic QSM visible in figure 7.

**FIGURE 7.**
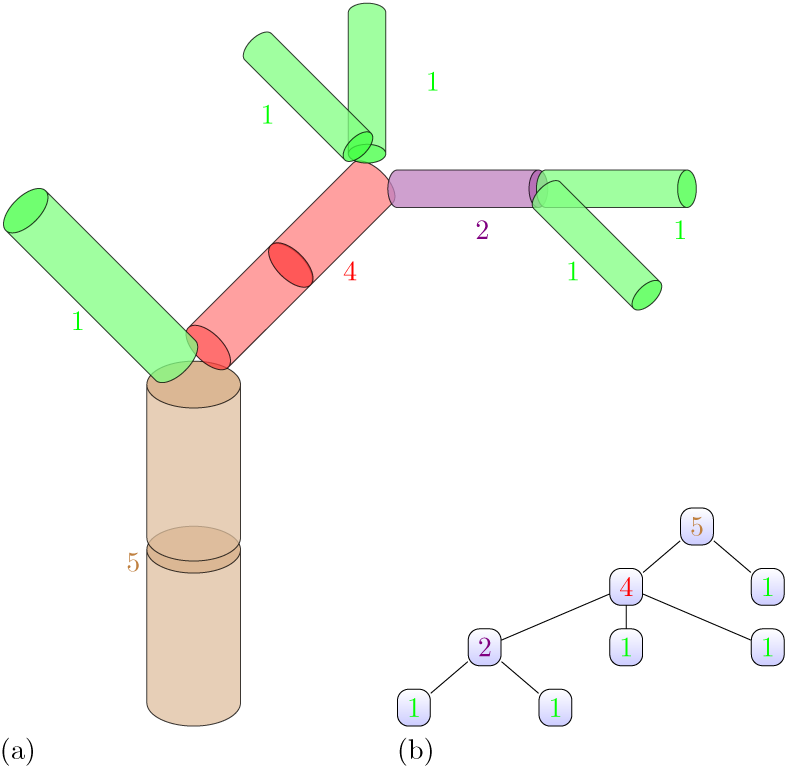
(a) The artificial QSM of figure 3 labeled with the `RPABO`. (b) We can see the topological order information from the QSM in (a) and each node has the `RPABO` displayed.

We conclude that the `RPABO` is the softwood area measured in an unknown metrical unit.

### 2.4 Reverse Pipe Radius Branch Order

Earlier works have shown that small cylinders can represent the topology correctly, but their radius tends to be hugely overestimated. By having a radius independent proxy for the cross sectional area we can simply take its square-root to derive a proxy for the radius of a cylinder.

#### Reverse Pipe Radius Branchorder

We define the Reverse Pipe Radius Branchorder (`RPRBO`) as the square root of the `RPABO`.

## Application for the Reverse Pipe Radius Branch Order

We can see the proxy potential in a clear visible linear pattern in figure 6 (c). So we will build the `RPRBO` filter to correct `radii` with equation 4.

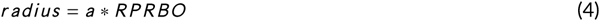

## 2.5 GrowthVolume

We defined in (Hackenberg et al., 2015a) the `GrowthVolume` of a cylinder recursively.

### GrowthVolume

The `GrowthVolume` of a cylinder is the cylinder’s volume plus the `GrowthVolumes` of its children.

In fact the `GrowthVolume` corresponds to the volume of a branch which is sawed from the real tree if we use the saw at the cylinder’s position. We used the `GrowthVolume` as a predictor for the radius in a radius correction filter, see 5.

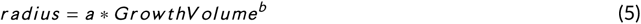

Nevertheless this filter has a design flaw. Overestimated `radii` of cylinders belonging to the `GrowthVolume` part effect their volumes and therefore also the `GrowthVolume`. So our predictor is effected by the noisy attribute we wanted to get rid of.

## 2.6 GrowthLength

We can substitute the volume in our previous definition to get a recursive defined length.

### GrowthLength

The `GrowthLength` of a cylinder is the cylinder’s volume plus the `GrowthLength` of its children.

## Application for the GrowthLength

We tested the `GrowthLength` also as a radius predictor, but results have not been really convincingly. But we can use it as the ordering predicate which we discuss in figure 3. The segment with the largest `GrowthLength` is always the first child segment.

### 2.7 ProxyVolume

**ProxyVolume** We can then further adapt to calculate the `ProxyVolume` of a cylinder by utilizing only the cylinder length and the `RPRBO` as shown in equation 6. By definition the `ProxyVolume` is not relying on correctly modeled `radii`.

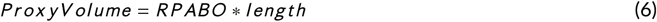

In previous works (Hackenberg et al., 2015a) we had defined the `GrowthVolume` of a cylinder in a recursive manner. We used the `GrowthVolume` as a predictor for a corrected radius of each cylinder utilizing a simple power function fit. Yet later produced SimpleForest results showed, that the influence of overestimated radius of twig cylinders lead to a too large overestimation of the `GrowthVolume` predictor. The radius correction filter therefore did not perform that well on unpublished user data.

We can though define the `VesselVolume` in similar manner, but utilizing instead of cylinders’ volumes their `ProxyVolumes`.

## 2.8 VesselVolume

### VesselVolume

The `VesselVolume` of a cylinder is its `ProxyVolume` plus the `VesselVolume` of the cylinder’s children.

## Application of the VesselVolume

As with the `GrowthVolume` we can try to predict the radius from the `VesselVolume` by a simple non linear power fit as we see in figure 6 (d). So we will use equation 7 to correct cylinders `radii` and will refer to as the `VesselVolume` filter.

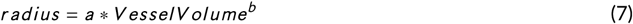

## 3 RESULTS

Point cloud data of 65 clouds and their calculated QSM Data produced by TreeQSM was used as pubished at Zenodo (Demol et al., 2021b). For TreeQSM models a tapering filter (Calders et al., 2015) to correct overestimated leaves was applied.

We compete against the accuracy of TreeQSM with our new implementation. In figure 8 we use SimpleForest QSMs which have been improved by either the linear `RBRBO` filter (a) or the non-linear `VesselVolume` filter (b).

**FIGURE 8.**
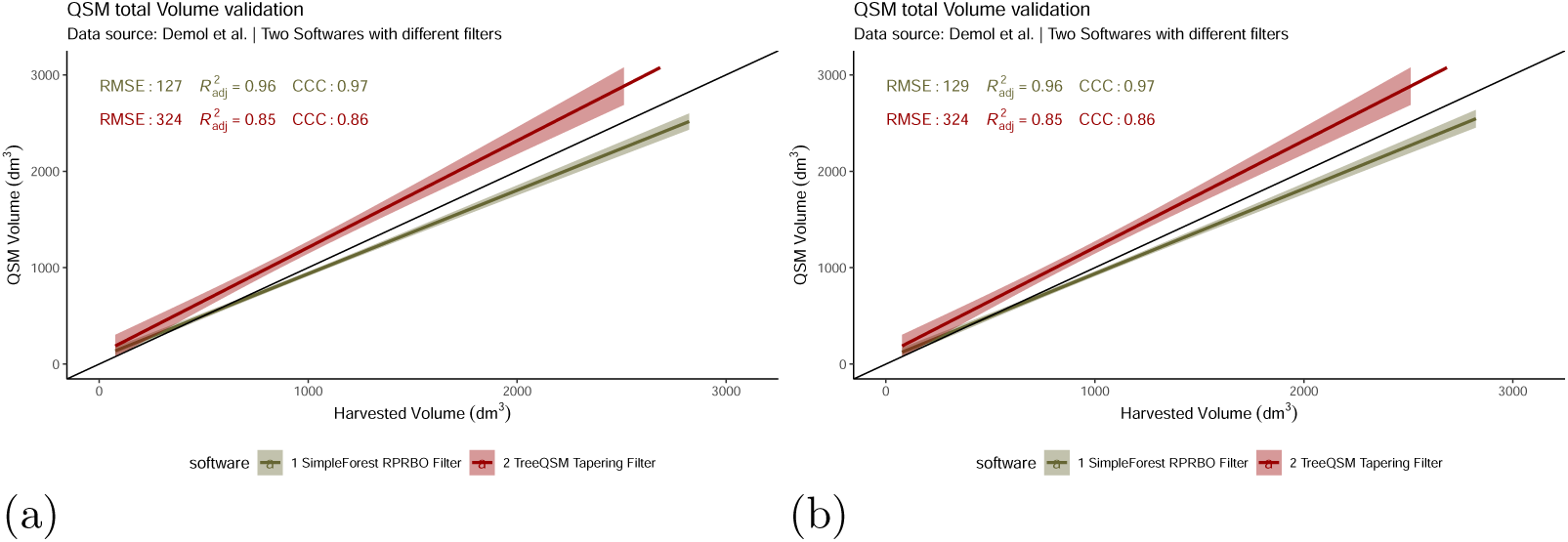
(a) The `RPRBO` filter performs well. The 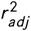. as well as the `CCC`are lot higher than those reached by The TreeQSM tapering filter. (b) The `VesselVolume` filter has accuracy of the same order.

Both results are written with their `RMSE`, their r^2^_adj._ and their `CCC` into table 1.

**TABLE 1.** Numerical validation of QSM volume measures compared to harvested reference volume measures.

| Software | RMSE ($dm^3$) | $r^2_{adj}$ | CCC |
| --- | --- | --- | --- |
| SimpleForest VesselVolume | 129 | 0.96 | 0.97 |
| SimpleForest RPRBO | 127 | 0.96 | 0.97 |
| TreeQSM Tapering | 324 | 0.85 | 0.86 |

## 4 DISCUSSION

We imported here in our analysis results (Demol et al., 2021a,b) which have been produced with TreeQSM. The TreeQSM pipeline which was utilized by the authors uses the algorithm described in detail (Raumonen et al., 2013) combined with optimization techniques discussed in (Hackenberg et al., 2015a) to achieve parameter adaptation. The QSMs then have been filtered with the tapering method (Calders et al., 2015). Importing the same clouds we also produced SimpleForest results relying on the same parameter optimization technique but utilizing a different algorithm (Hackenberg et al., 2014). And filtered with the here proposed allometric theory based filters. Our results produced data where the `RMSE` is 39% of the TreeQSM error. The devation of the r^2^_adj._ value from the best score 1.0 is for SimpleForest only 27% of the TreeQSM derived deviation. For the `CCC` the deviation is only 21% of the TreeQSM deviation. SimpleForest utilizing the here presented filters performs astonishingly well compared to the broadly accepted TreeQSM tool.

To get more insight into the error we want to split up the total error into a measurement error (error_ref_) of the destructively collected reference data, the error produced by the algorithm (error_alg_) which is limited by the potential of the filter to reduce exactly this error (potential_fil_), see equation 8.

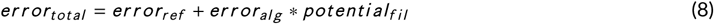

The error_ref_ is the exact same for TreeQSM as well as SimpleForest results. Therefore we will purely focus in the following on error_alg_ and potential_fil_. With the data here used we can’t show not more than SimpleForest with one of the allometric scaling filters performs better than TreeQSM with the tapering filters. For splitting up the error numerically a more complex result matrix would have to be generated. While out of scope for this work we consider the following analysis as quite useful. Produce on a preferably larger data set SimpleForest as well as TreeQSM and 3dForest results and then applying for each of the three softwares all of the three filters, e.g. produce in total nine results.

While the analyzed data is not sufficient large enough to be able to perform a prove with the presented result table, we do the hypothesis that error_alg_ has much less impact than potential_fil_. As we developed not only the filter, but also the whole SimpleForest plugin we have a lot of expert knowledge. Earlier collaborations with the researcher behind TreeQSM (Hackenberg et al., 2015a) as well as pre-printed SimpleForest software results under rework (Hackenberg et al., 2021) give a link that the accuracy of the algorithm is less impactful. Whenever we used other filtering techniques with SimpleForest we reached worse results with an error of approximately the magnitude of TreeQSM’s error.

While we still have to do more numerical proofs in the future we can conclude to some extend that the analyzed tree species can be modeled well with the here presented allometric scaling theory based filters. We consider the error measures too good to be random and as we fit a function with only two parameters over-fitting potential is rather low.

Excluding the aspect of robustness the formula given in equation 7 is not be the best choice. If we look more carefully into the tree data we can think of the `VesselVolume` radius relation as a temporal growth function. See figure 9 for a scatter plot of one QSMs where the unfiltered QSM already revealed good fitting quality, we swaped here x and y axis.

**FIGURE 9.**
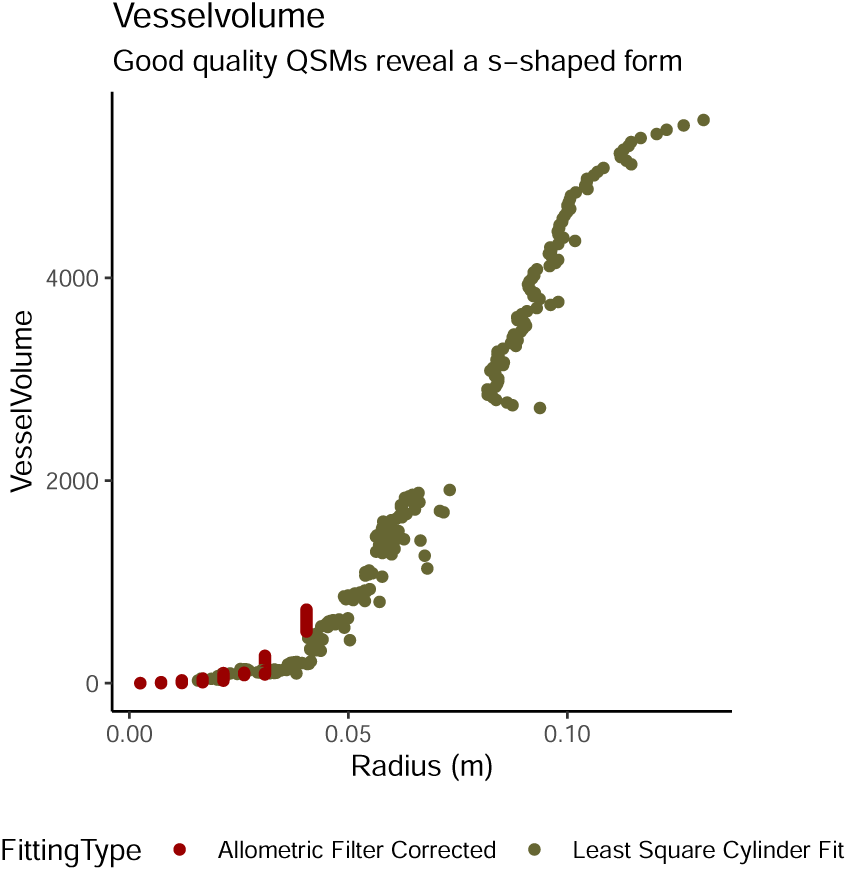
We see cylinder measurements of one QSM with good quality in our data set. The green dots correspond to unmodified cylinders and the red to ones which have been filtered with the here `VesselVolume` technique. Due to the good quality nature of the analyzed QSM we see a plateau effect. The green part of the curve flattens for the largest cylinders, the trunc cylinders. The whole curve can be described with a s-shaped five parameter based temporal growth function.

The s-shaped form would obviously still be visible if we exchange predictor and predicted but in a rotated manner. So we might think of applying inverted growth function utilizing `VesselVolume` and `radius`. Commonly used non linear s-shaped functions contain five parameters and are harder to implement in a robust way inside a distributed software. If data gets more noisy figure 9 depicts only data from a well fitted QSM - fitting five parameters is supposed to sometimes fail. Still we hope that we cause with our work a lot of followup works also utilizing growth functions. Implementing a decent prototyped filter into C++ to give convenient access to SimpleForest users can then later be done. SimpleForest allows the embedding of citable work. As our function ignores the upper right plateau which visible in figure 9, the largest `radii` can be underestimated by our power function `VesselVolume` filter. We can note that we face underestimation in our data and this might be at least partially be solved by utilizing another model as proposed.

We conclude that the applied filter shows that the combination of the PMT and the WBE model give a lot of insight and good modeling capabilities for the branching structure of tree. But unexplained is the following fact. We already stated in the definition section that the `RPABO` equals the number of supported twigs - cylinders with a recursive child relation to the query cylinder but no own children. The PMT states that this is the area of the softwood vessel at the query intersection. But we fit the total `radius`, which includes also the inner hardwood. And as the radius can be converted to an area via the simple geometry circle formula our predictor can be seen as the area of soft and hardwood.

But it is a bit strange that we can predict from a pure softwood area only the combined soft and hard wood area. This works out if the tree had in the last growth season the average expected growth potential. Trees yet tend to growth less than expected in hot and dry years. In such years less softwood is expected and we would expect a bias here in the total wood area prediction. This is of course also a second explanatory hypothesis for the bias we see in the SimpleForest result plots.

If this hypothesis would get strengthened we would have to increase the model complexity by having a one growing season long climate proxy. Climate data draw spatial temporal patterns. Future better models might have to include many parallel geo-tagged time-series.

## 5 CONCLUSIONS

QSMs contain a lot of measurements. The allometric scaling theory can also applied to the scaling of other species and also not only to trees (Packard, 2020). Even mammals can be model in such manner. Research as the cited one for the broad field of biology can benefit as well to get insight into allometric scaling by utilizing trees and their QSMs as example data to build a generic theory onto. Data for trees is rather easy to collect as we have non-destructive measuring techniques applied on plants which in contrast to animals don’t change location over time.

The impact of QSMs to forest ecology theories should hopefully be clear. While collecting TLS data still requires access to high tech devices we also mentioned in the introduction, that the point clouds of tomorrow can be derived from today’s photographies. Tech giants like Google/Alphabet or Facebook/Meta can be considered having an oligopol on such data. It is therefore from a huge importance to provide the world with free, open and scientific data bases as well to build here an academical counter part to the industry dominance. A well designed ecological smart phone app with a huge data base server as backup is doable with some efforts and fundings. User accounts should require optionally the OrcID to be able to separate academical users from non-academical personnel. We read news paper articles about other eco system related projects where people count birds as a hobby and feed their results into prepared data bases. Such approach can also happen in forestry, but it requires maintenance and setup work.

### 5.1 Future Work

We first want rework the preprinted SimpleForest software paper to provide an up to date reference for the here used free software tool (Hackenberg et al., 2021).

We also identified the need for a data paper. Various data sets have already been made accessible (Demol et al., 2021a; Hackenberg et al., 2015b; de Tanago et al., 2018) to combine all small data bases to a larger one with backuped by a manuscript might convince more data owners to share their data and instead of 65 trees proposed future analysis can be run on few hundred of tree clouds with harvested volume data back-uped.

## acknowledgements

Acknowledgements should include contributions from anyone who does not meet the criteria for authorship (for example, to recognize contributions from people who provided technical help, collation of data, writing assistance, acquisition of funding, or a department chairperson who provided general support), as well as any funding or other support information.

## Supporting Information

Supporting information is information that is not essential to the article, but provides greater depth and background. It is hosted online and appears without editing or typesetting. It may include tables, figures, videos, datasets, etc. More information can be found in the journal’s author guidelines or at http://www.wileyauthors.com/suppinfoFAQs. Note: if data, scripts, or other artefacts used to generate the analyses presented in the paper are available via a publicly available data repository, authors should include a reference to the location of the material within their paper.

---

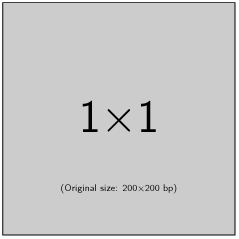

**A. One** Please check with the journal’s author guidelines whether author biographies are required. They are usually only included for review-type articles, and typically require photos and brief biographies (up to 75 words) for each author.

## GRAPHICAL ABSTRACT

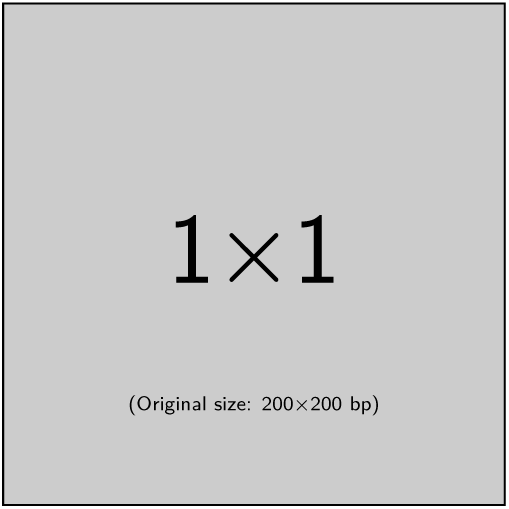

Please check the journal’s author guildines for whether a graphical abstract, key points, new findings, or other items are required for display in the Table of Contents.

